# How SARS-CoV-2 alters the regulation of gene expression in infected cells^†^

**DOI:** 10.1101/2022.12.18.520908

**Authors:** Emmanuelle Bignon, Stéphanie Grandemange, Elise Dumont, Antonio Monari

**Author notes:** corresponding authors: E.B, E.D., A.M. Electronic Supplementary Information (ESI) available: Time series of the RMSD for all the systems, time series of the crucial distances between ARKS and the closest nucleotides, binding free energies between ORF8 and DNA also in presence of acetylation/methylation of K36.

## Abstract

Non-structural accessory proteins in viruses play a key role in hijacking the basic cellular mechanisms, which is essential to promote the virus survival and evasion of the immune system. The immonuglobulin-like open reading frame 8 (ORF8) protein expressed by SARS-CoV-2 accumulates in the nucleus and may influence the regulation of the gene expression in infected cells. In this contribution, by using micro-second time-scale all-atom molecular dynamics simulations, we unravel the structural bases behind the epigenetic action of ORF8. In particular, we highlight how the protein is able to form stable aggregates with DNA through a histone tail-like motif, and how this interaction is influenced by post-translational modifications, such as acetylation and methylation, which are known epigenetic markers in histones. Our work not only clarifies the molecular mechanisms behind the perturbation of the epigenetic regulation caused by the viral infection, but also offers an unusual perspective which may foster the development of original antivirals.

## Introduction

Since its first outbreak at the end of 2019, and the declaration of pandemic state in Spring 2020, COVID-19, and its causative agents, the RNA positive-strand beta-coronavirus SARS-CoV-2 have been the subject of intense studies, clarifying many fascinating cross-talks between the viral and cellular machinery.^1,2^ In many cases, such interactions have also highlighted viral adaptative strategies allowing its evasion of the immune system response, or increasing its transmissibility helping spreading the infection further.^3,4^ Even if after more than two years, the COVID-19 epidemic may be considered under relative control, mostly thank to the development and the deployment of novel vaccinal strategies, including mRNA agents,^5,6^ the necessity to precisely understand the complex viral machinery and its interplay with cellular regulation still holds. This is even more stringent, since RNA viruses are among the most threatening emerging pathogens,^7–9^ due to their high mutation rate, easy and widespread diffusion, and to the facile interspecies barrier crossing. Therefore, biophysical and biochemical approaches, coupled with cellular and molecular biology, are still needed to clarifying different viral mechanisms, including resolving the structure of a number of viral accessories proteins, and elucidating their biological function.^10–12^ Indeed, upon infection the genome of SARS-CoV-2 is directly translated, by the cellular machinery, into structural and non-structural proteins (Nsp). While the former, including the well celebrated Spike protein, are incorporated into the nascent virions, the latter either perform crucial viral processes to assure the replication, or interact with cellular partners to hijack and perturb the cellular processes. Some of the most well-known Nsp include the papain-like and main protease,^13–15^ which cleaves the unfunctional viral polyprotein into functional units, and the RNA-dependent-RNA-polymerase,^16–18^ which replicates the viral genome. Because of their importance in the virus life-cycle, and particularly in replication and maturation, these Nsp are targets of choice for the possible development of antiviral drugs, including the, up-to-date, only commercialized anti SARS-CoV-2 agent, which is a covalent inhibitor of the main protease.^19^

In addition to those exerting an enzymatic activity, other Nsps exist which perturb the cellular machinery, notably limiting the global immune response of the cells.^20^ As an example, this include units sequestering proapoptotic cellular RNA like the SARS Unique Domain,^21,22^ as well as membrane proteins influencing autophagy (Nsp6)^23,24^ or acting as porins to permeabilize and depolarize the cellular membrane (Open Reading Frame 3, ORF3).^16,25–27^

Recently, the role of ORF8 has been particularly pointed out.^28– 30^ Indeed, this small water-soluble immunoglobulin-(Ig)-like protein is interfering with the immune system response, particularly by interacting with monocytes.^31^ The structure of ORF8 in bat and human coronaviruses has also been resolved, and the molecular bases of its perturbation of the immune response, which proceeds through the formation of sulphur-bridge mediated large scale aggregates, has been discussed.^30,32^

However, the importance of ORF8 goes way beyond this direct Ig-like effect. More recently, it has been shown that ORF8 is able to penetrate the nuclear membrane and accumulates in the nucleus.^33^ Furthermore, ORF8 has been co-precipitated with chromatin proteins and has been shown to be subjected to histone-like post-translational modifications (PTMs), such as lysine acetylation and methylation.^33^ Finally, the presence of ORF8, and its post-translational modification, has been associated to the perturbation of the chromatin compaction, and to the alteration of the epigenetic profiles of the infected cells (see Figure 1).^33^

**Figure 1.**
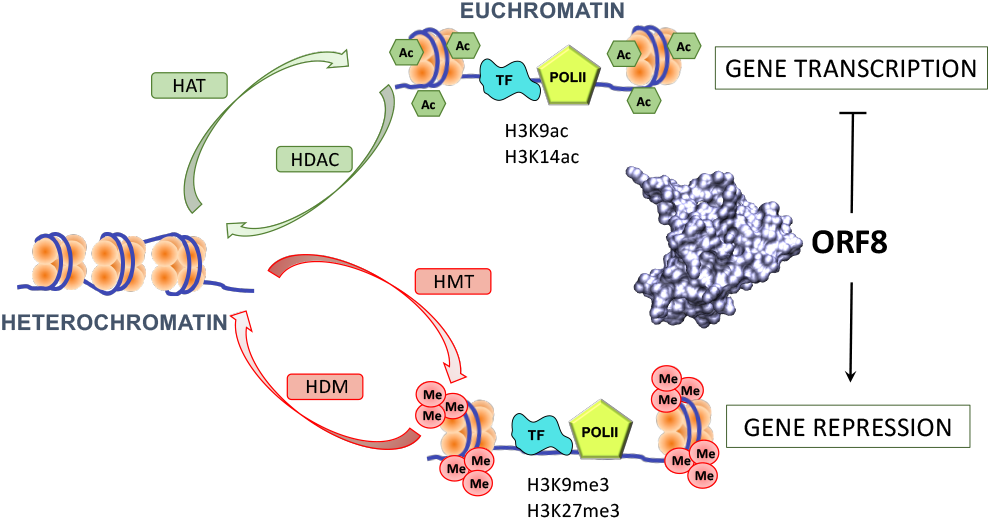
Exemplification of the action of ORF8 in promoting chromatin phase-transition and hence modulating the epigenetic responses in the infected cells. The role of PTMs, and in particular the ORF8 induced perturbation of the H3 histone acetylation (HAT)/deacetylation (HDAC) leading to the repression of gene expressions is particularly evidenced.

Thus, ORF8, is behaving as a histone mimicry, and influencing the euchromatin/heterochromatin transition is also clearly acting as an epigenetic regulator. If the ensemble of data confirming this action is firmly rooted, the knowledge of the molecular and structural bases of this regulation is still rather scarce. In particular, no structure of the complex between ORF8 and DNA or chromatin proteins has been presently resolved. In this contribution we aim at filling this gap by using a multiscale approach combining protein/nucleic acid docking and μs-scale all atom-molecular dynamic (MD) simulations to present the first model of the complex between ORF8 and double-helical DNA. The obtained stable and persistent aggregates are coherent with the observed coprecipitation of ORF8 with chromatin. Furthermore, we also evidence how PTMs modify the interaction network between DNA and ORF8 and hence are susceptible to induce chromatin transitions. To the best of our knowledge, our contribution is the first study detailing, at atomistic level, the effects of viral infections into the deregulation of gene expression and epigenetic profiles.

## Results and Discussion

From a structural point of view29 ORF8 is a rather small protein (106 aminoacids) presenting a high density of μ-sheets, which provide a rigid core, decorated with rather flexible and unstructured loops. ORF8 crystallizes in the form of a dimer (Figure 2), while the supra molecular organization may be essential for the Ig-like action, the monomer form is also considered as functional, especially for its interaction with DNA and chromatin. The global rigidity of the core is also reinforced by the presence of three disulphide bridges per monomer. An additional intermolecular sulphur bridge at the protein/protein interface is present. The analysis of the MD simulation of ORF8 in its monomeric and dimeric form confirms the stability of both systems, as also shown by the relatively small and stable evolution of the root mean square deviation (RMSD) as shown in Supplementary Information (SI). Interestingly, the effects of the intermolecular sulphur bridge seem rather marginal on the global stability of the dimer, as shown by the similar evolution of the distance between the centres of mass of the two monomers in the bridged and unbridged system as also reported in SI.

**Figure 2.**
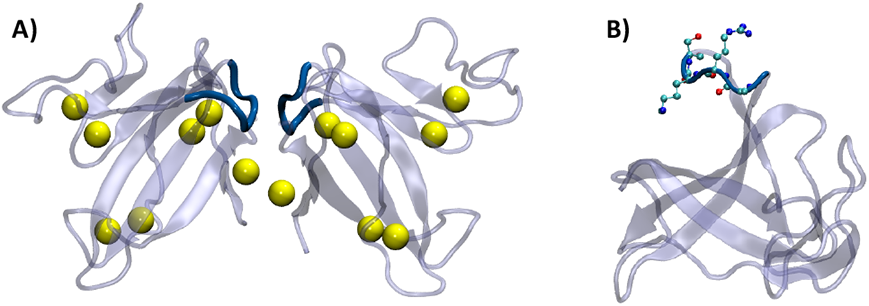
Representative snapshots issued from the MD simulations for the apo ORF8 in dimeric (A) and monomeric (B) form. The possible disulphide bridges are evidenced in A) by showing the position of the interacting sulphur atoms. The position of the ARKS motif is evidenced in blue colour and the corresponding aminoacids are explicitly represented in panel B.

Of particular importance is the presence, on one of the flexible loops, of the ARKS sequence (residue 35 to 38) and to a lesser extent of the subsequent AP motif (residue 39 and 40). Indeed, this motif, which is highlighted in Figure 2 presents a very strong sequence similarity with the one found in the flexible tail of the histone 3 (H3).^33^ Besides, lysine 36 is also considered a hot-spot for PTMs,^34–36^ which may lead to different outcomes relative to gene expression or silencing. Because of the sequence similarity, and also considering the high density of positive charges, the ARKS sequence may clearly be regarded as one of the leading factors favouring the interaction with DNA, and thus to a more global level chromatin compaction. Interestingly, and as can be assessed from Figure 2, the ARKS motif is highly exposed in the monomer, while is relatively buried at the protein/protein interface in the case of the dimer, and hence less accessible for DNA complexation. This fact, also confirms the hypothesis that the monomeric form of ORF8 should be active towards DNA.^33^ To confirm the possible formation of stable aggregates between ORF8 and DNA we have performed protein/nucleic acid docking to obtain starting conformations. Two poses have emerged from the procedure as the most stable or mostly populated cluster and are reported in SI. The two poses have further been relaxed through μs-scale all atom MD simulations. The MD has confirmed that ORF8 is interacting persistently with DNA strands, as shown from the representative structures reported in Figure3. However, the first docking pose, which already provided DNA contacts with the ARKS motif evolves only slightly and leads to the insertion of the ORF8 tail in the DNA major groove. Instead, the second pose, initially presented the ARKS tail oriented away from the duplex. However, upon MD and on the hundreds of ns time-scale ORF8 relaxes and slightly slides along the duplex, to finally stabilizing after establishing a contact between ARKS and the nucleic acid minor groove (Figure 3B).

**Figure 3.**
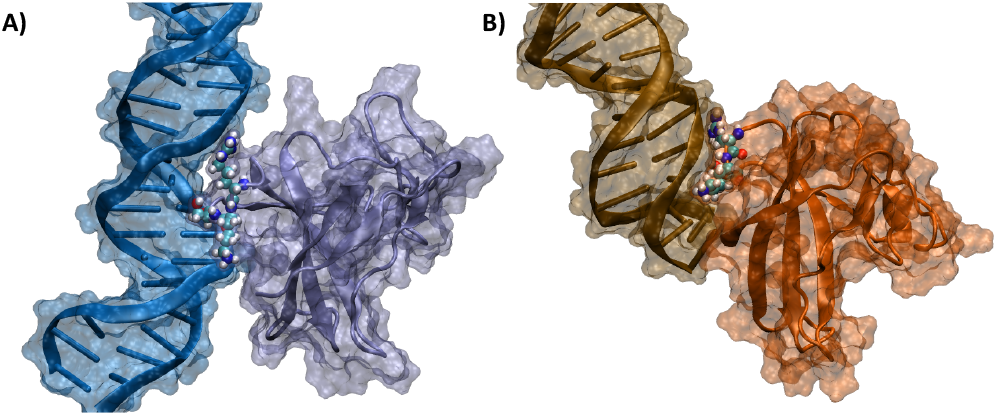
Representative snapshots of the two identified interaction modes of ORF8 with a DNA strand, namely with major (A) and minor groove (B). The ARKS motif is evidenced in van der Waals representations.

Although globally stable, the two interaction modes give rise to slightly different interaction patterns involving, mainly, the nucleic acid backbone. Notably, however, both the major and the minor groove bound states involve the ORF8 RKS sequence which develops rather persistent interaction with the nucleic acid fragments. Globally, these results are also coherent with the experimental observation of the knocking-off of the interaction with chromatin by deletion of the ARKS sequence in ORF8.^33^

In Figure 4 we report the distribution of the evolution of crucial distances between the RKS aminoacids and the closest nucleotide, the corresponding time-series are also provided in SI. As concerns R36, the interactions mainly involve the backbone and the phosphate moieties, as it is expected due to the complementary charges. Indeed, we may evidence a rather strong interaction between the guanidino group of the aminoacid and the phosphate group of T11 leading to an evident peak at a distance of about 1.8 Å which is compatible with hydrogen bond (HB) formation. Although, the distribution appears quite peaked, indicating a rather dominant interaction, the analysis of the time-series shows that this HB is, indeed, more mobile and multiple, temporary and short lived, occurrences of conformation with higher distances are populated. The temporary breaking of the interaction is due to the slight rotation of the aminoacid away from the nucleotide. However, this movement may approach the guanidino group of R36 to the phosphate moiety of thymine T12. As shown in the second panel of Figure 4, a bimodal distribution is observed for this distance. A rather sharp peak at around 1.8 Å, and a shallow, band with maximum at around 4.0 Å. Therefore, we may conclude that a secondary interaction takes place with this nucleotide, compensating for the temporary loss of the interaction with T11 and globally reinforcing the stability of the protein nucleic acid aggregate. The rather non-persistent nature of this interaction may however be appreciated from its time-series (see SI) which shows strong oscillations between bound and unbound states.

**Figure 4.**
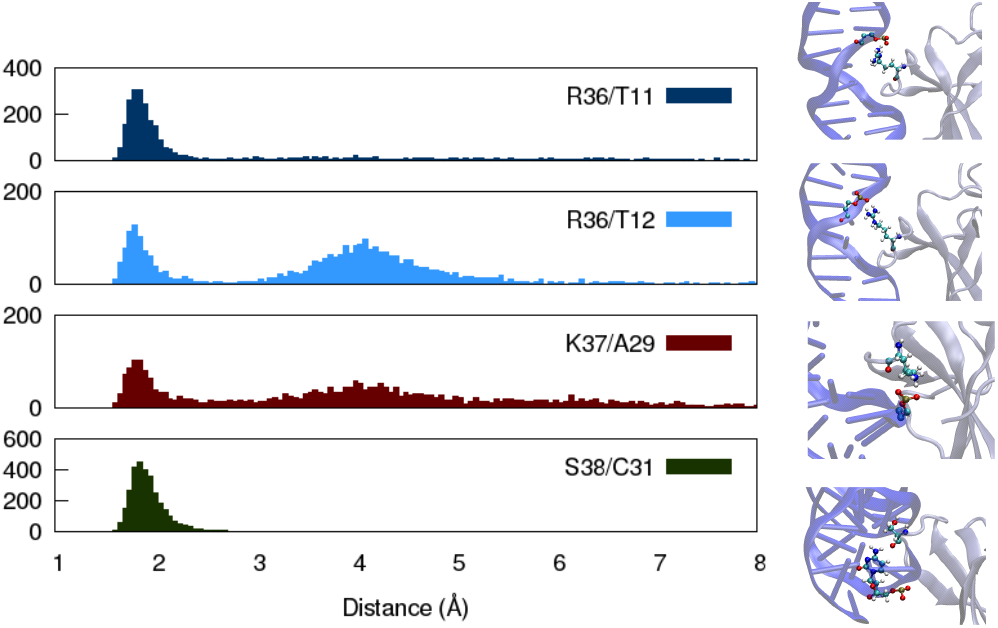
Distribution of the distances between the R36, K37, and S38 aminoacids and the most proximal nucleotides for the major-groove-bound ORF8. The distances are calculated between the closest OP atom of the nucleotide and the guanidino hydrogen of R36 and the amine hydrogen of K37. The distance between the amine group of Cytosine and the carbonyl oxygen of S38 is also reported. Representative snapshots of the interactive units are also provided.

Unsurprisingly, the main interactions developed by K37 involves the nucleic acid backbone and most specifically the phosphate group of A29 and the charged amino group of this aminoacid. However, it is also evident that the interaction of K37 is less pronounced than for R36. The distribution of the distances still presents a rather sharp peak around 1.8 Å, however, the latter is weaker than for R36 and the distances are definitively more spread towards larger values. Furthermore, a bimodal distribution is also clearly evident with a secondary peak, belonging to a very large and shallow band, appearing again at 4.0 Å. Differently, from the case of R36, the lability of the interactions of K37 is not compensated by the development of secondary interaction patterns with other nucleotide, whose distances from the aminoacid charged amino group never decrease below 5.0 Å.

Finally, we may instead observe (Figure 4 bottom panel) a very strong and persistent interaction between S38 and C31. If the involvement of this aminoacid in the recognition of DNA has already been hypothesized, rather surprisingly it involves the HB between the carbonyl oxygen of the protein and the amine group of cytosine C31. However, the interaction is extremely persistent and leads to a very peaked maximum at 1.8 Å comprising almost all the occurrences. This is also evident in the time-series reported in SI in which no loss of the interaction could be seen during the whole span of the MD simulation exceeding the μs-scale.

The main interactions at the residue level for the minor groove-bound ORF8 are instead reported in Figure 5, while the corresponding time-series are provided in SI.

**Figure 5.**
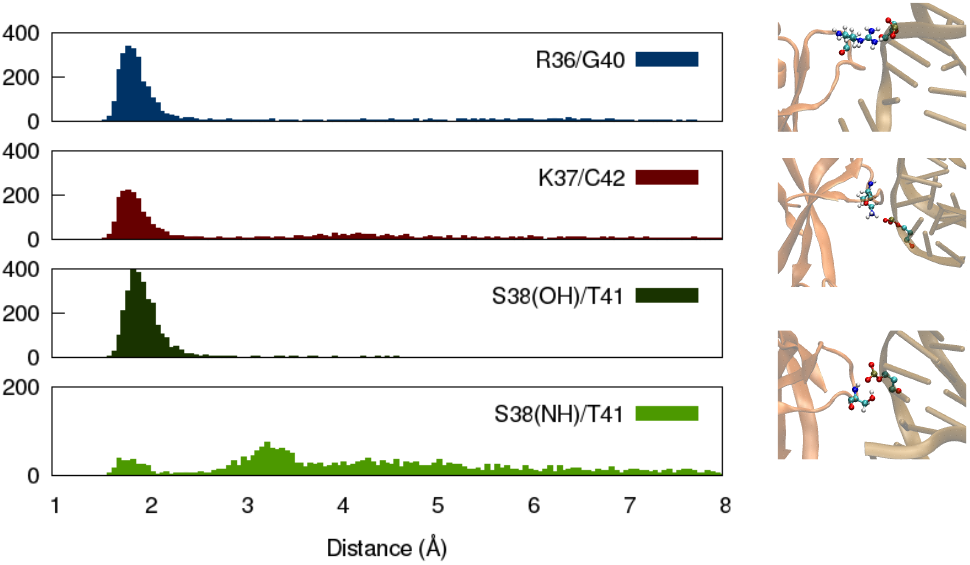
Distribution of the distances between the R36, K37, and S38 aminoacids and the most proximal nucleotides for the minor-groove-bound ORF8. The distances are calculated between the closest OP atom of the nucleotide and the guanidino hydrogen of R36, the amine hydrogen of K37, as well as the lateral chain hydrogen of S38, and the amino hydrogen of the backbone of S38. Representative snapshots of the interactive units are also provided.

In the case of R36 we may now observe a well-defined peak at around 1.8 Å for the distance between the hydrogen of the guanidino group and the closest oxygen of the phosphate moiety. While this specific interaction appears more persistent than the corresponding one for the major groove-bound conformation, no secondary interactions of R36 with other nucleotides can be evidenced from the MD simulation. Interestingly, the persistence of the interaction of K37 with the phosphate oxygen is also maintained in the minor groove-bound state, and the bimodal character of the interaction, which was evident for the major groove-binding, has now almost vanished and only the principal peak at 1.8 Å survives. Coherently, oscillations between bound and unbound states in the time series are also damped noticeably. Finally, as concerns S38, its interaction with DNA is reverted to a more classical pattern involving the hydroxyl group of the lateral chain and the phosphate backbone. Again, the corresponding HB is stable and highly persistent all along the MD trajectory as shown by the peaked distribution and by the very moderate oscillations experienced by the time-series. Interestingly, a secondary interaction between the backbone -NH group of S38 and the phosphate moieties can also be evidenced (see bottom panel of Figure 5). While this interaction is much less persistent and highly labile, a small peak at distances compatible with HB formation may be evidenced. Thus, it can participate in the global stabilization of the protein/nucleic acid complex. The estimation of the binding free energies however highlights significant differences between the two poses. As a matter of fact, the major-groove bound state appears favourable leading to a binding free energy of -45.6±8.2 kcal/mol. Conversely, the minor groove binding only provides a stabilization of -28.4±9.6 kcal/mol.

Globally, and coherently with the experimental observations, our results prove that ORF8 is indeed able to stably bind DNA via its histone-mimicry ARKS motif. More specifically, two motifs have been evidenced involving either the DNA major or minor groove Whether, the second one is probably less favourable, both motifs may be competitive to assure DNA binding. Therefore, ORF8 is susceptible of influencing the compaction of the chromatin and hence to act as a perturbator of the epigenetic regulations of the host cells.

The ARKS sequence in the H3 histone being also a hotspot for PTMs, and particularly lysine acetylation and methylation, in the following we will examine their effects on the stability of the ORF8/DNA aggregate and on the modification of the established interaction network.

Even upon the presence of PTMS, and more specifically lysine acetylation or methylation, the ORF8/DNA complex remained stable throughout the MD simulation, without showing any relevant structural modification either in both interaction modes. Unsurprisingly, however, the interaction of K37 with the DNA backbone is lost, whatever the PTM considered as shown from the distance distribution reported in Figure 6.

**Figure 6.**
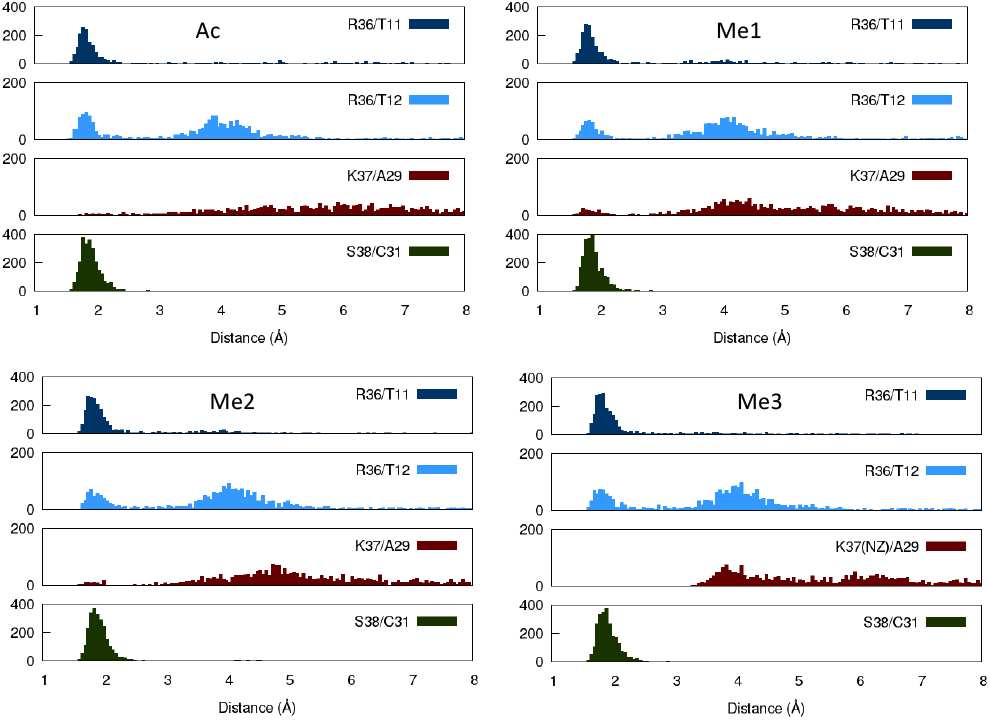
Distribution of the distances between the R36, K37, and S38 aminoacids and the most proximal nucleotides for the major-groove-bound ORF8 in presence of PTMs on K37 (Ac: acetylation, Me1 single-methylation, Me2 double-methylation and Me3 triple methylation). The distances are calculated between the closest OP atom of the nucleotide and the guanidino hydrogen of R36, the amine hydrogen of K37, as well as the lateral chain hydrogen of S38, and the amino hydrogen of the backbone of S38, with the exception of Me3, for which the distance to NZ is reported

This loss of interaction, which could involve chromatin remodelling, and favour gene expression, is however not accompanied by a noticeable weakening of the contacts experienced by the ARKS motif with DNA. Indeed, both the interaction of R36 and S38 are highly conserved, as shown in Figure 6 for the more favourable major groove interaction mode and Figure 7 for the minor groove one. Indeed, only rather negligible differences in the distribution of the distances compared to the native protein are evidenced in both the position and the shape of the main peak. Globally a more pronounced destabilization of the interaction network can be observed for the minor-groove bound mode, which is particularly evident for the acetylated and single-methylated ORF8 (Figure 7). Notably, in the latter case the more important perturbation are observed in the case of acetylation or single methylation, which is also coherent with the hypothesis of the histone code.^37^ The absence of a noticeable destabilization of the ORF8/DNA aggregate upon the inclusion of PTMs is also confirmed by the values of the binding free energy, which are reported in SI and do not show significant differences for the native or the acetylated/methylated ORF8.

**Figure 7.**
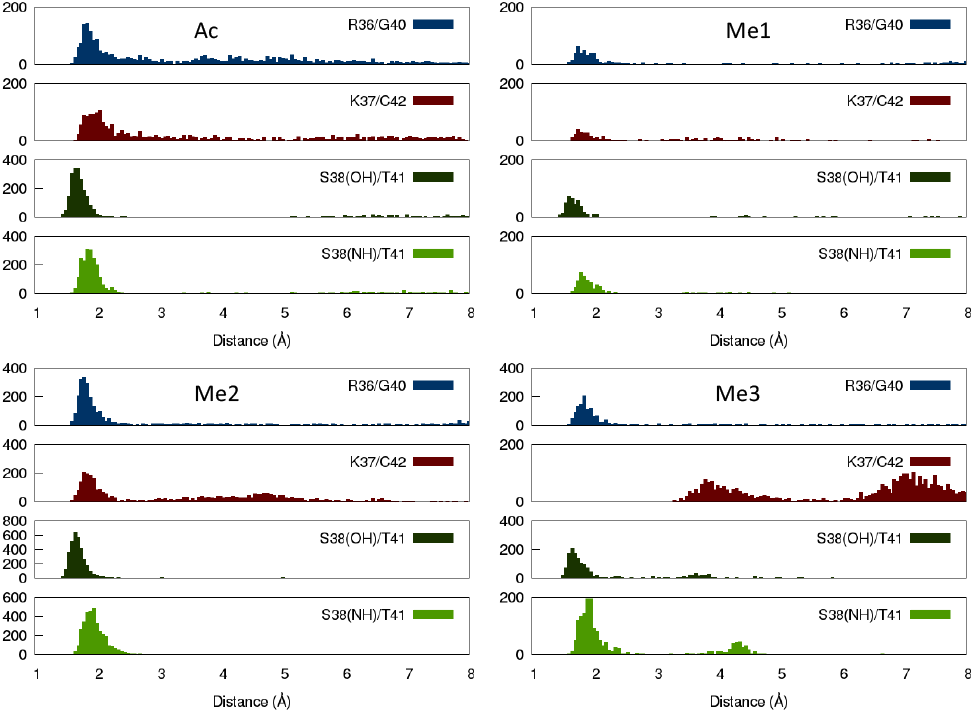
Distribution of the distances between the R36, K37, and S38 aminoacids and the most proximal nucleotides for the minor-groove-bound ORF8 in presence of PTMs on K37 (Ac: acetylation, Me1 single-methylation, Me2 double-methylation and Me3 triple methylation). The distances are calculated between the closest OP atom of the nucleotide and the guanidino hydrogen of R36, the amine hydrogen of K37, as well as the lateral chain hydrogen of S38, and the amino hydrogen of the backbone of S38, with the exception of Me3, for which the distance to NZ is reported

Globally our results confirm the experimental evidences which reports that cells infected with viruses expressing ORF8 present a global more compact chromatin and, thus, an important under expression of various genes, especially those involved in the immune response.^33^ Additionally, while our findings are coherent with the role of histone mimic of ORF8, they can also help interpreting the interplay with PTMs. Indeed, we have shown that ORF8 ARKS motif is able to maintain persistent interactions with DNA even in presence of methylation and acetylation. It can therefore be surmised that, while ORF8 may effectively competes with the native H3 as a substrate for PTMs-inducing enzymes, the introduction of acetylation and methylation on the viral protein will induce a different biological effect. Indeed, the presence of PTMs on ORF8 will not result in chromatin decompaction, hence it will perturb the epigenetic profile of the infected cell and thus favour the viral reproduction. However, to confirm this hypothesis the interaction between ORF8 and acetylation and methylation enzymes should be properly modelled and simulated.

## Conclusions

By using full atom molecular simulations, we have unravelled the behaviour of a SARS-CoV-2 accessory protein namely ORF8. In particular, we have confirmed that ORF8 is indeed able to interact with double helical DNA, through its ARKS motif, which is reminiscent of the histone tails. Therefore, and since ORF8 may bypass the nuclear membrane and accumulate in the cellular nucleus, it can influence the chromatin compaction of the host cell, and ultimately altering the gene expression profile of the infected organisms.

We have evidenced two stable interaction motifs of ORF8 with double strand DNA through either the major and the minor groove. In all cases stable interactions between arginine/lysine and the backbone phosphate are developed, while the serine is assisting the compaction through hydrogen bonds.

We have also evidenced that the inclusion of PTMs, such as acetylation and methylation on the lysine in the ARKS motif, maintain a stable aggregate with the DNA, and are not susceptible to strongly alter the interaction patterns involving arginine and serine. Thus, we believe that not only ORF8 may participate in the DNA compaction, but it can also compete with the native histone tail draining the H3-oriented modification aimed at favouring gene expression. Indeed, when introduced on ORF8 PTMs appears to have, coherently with the experimental results,^33^ only a moderate effect on the chromatin compaction, and hence in gene expression. Therefore, ORF8 may accumulate PTMs and divert their action neutralizing the cross-talks leading to chromatin remodelling, and, thus, avoiding the expression of genes potentially activating the immune response.

To the best of our knowledge, our study represents the first modelling of the epigenetic effects induced by a viral protein on the host cells. It helps in unravelling the complexes molecular mechanisms and cross-talks put in action by viruses to hijack the infected cell machinery and optimize their survival and reproduction.

In the near future we plan to also study the interaction between ORF8 and acetylation and methylation enzymes to assess the hypothesis of the facile accumulation of PTMs on the viral accessory protein. In this respect our results may also pave the way to original therapeutic strategies based on counteracting the chromatin compaction action of ORF8 to weaken the virus defences against the immune system response.

### Computational Methodology

The initial ORF8 structure has been retrieved from the one crystallized by Flower et al. (PDB 7JTL)^29^ the missing aminoacids have been reconstructed by homology model using the SwissModel web server.^38^ Sulfur bridges between the neighbouring cysteines have been enforced consistently during the MD simulation. Both the crystallized dimer and monomer have been simulated. In the case of the dimer we have considered both the situation in which the two monomers are bound via an interprotein sulfur bridge or the one in which only intermolecular interactions are present, showing that the intermolecular bridge has only a limited effect on the stability of the aggregate. The protein has been represented using the amberff14sb force field,^39^ the ARKS-containing loop is usually known to exhibit a high degree of disorder, hence we have included the grid-based energy correction maps (CMAP) describing intrinsically disordered proteins^40–43^ to the aminoacid 30 to 50 of both monomers. Each initial system has been solvated in a cubic box of TIP3P^44^ water molecules including a 9 Å buffer. After minimizations the systems have been thermalized and equilibrated by progressively releasing positional constrains on the protein backbone atoms during 9 millions time-step. The MD simulation has been performed in the isobaric and isothermal (NPT) ensemble using Hoover thermostat and Langevin barostat.^45^ The Newton equations of motion have been numerically integrated using a time-step of 4 fs, thanks to the use of the Hydrogen Mass Repartition (HMR)^46^ strategy in combination with the Rattle and Shake algorithm,^47^ and propagated for about 1.0 μs. Representative conformation of the apo ORF8 in both dimeric and monomeric state have been obtained by clustering of the trajectory based on the RMSD state. A 21 base pairs model B-DNA duplex of sequence 5’-CGGACATTCTTCCGGTTGGAC-3’ has been docked to both OPRF8 and ORF8 monomer using the HADDOCK web server.^48^ The stability of the docking poses have been further assessed by performing MD simulations of the protein/DNA complex. The same parameters as for the apo systems have been applied while DNA is modelled by amberff99sb force field including the bsc1 corrections.^49,50^ The docking to the dimer gave rise to a rather unstable complex, and does not allow to develop interactions between ARKS and the nucleic acid, and is therefore further discarded. Conversely, the poses for the monomer yielded stable structures and are therefore further analysed, with simulations exceeding 1.2 μs.

To take into account the effects of the PTMs on K36 we have in silico modified this aminoacid introducing one acetyl group, as well as one (Me1), two (Me2), or three (Me3) methyl moieties on the lateral chain amino group. Each PTMs has been introduced on representative snapshots issued from the two interaction modes, obtained from a clustering of the MD simulations, thus resulting in 8 independent initial systems. For each of the K36-modified conformations we have again run equilibrium MD simulation, using the same protocols as for the native complex, for a total time of 1.0 μs.

All the equilibrium MD simulations have been performed using the NAMD code,^51,52^ and the trajectories have been visualized and analysed through the VMD^53^ and *ccptraj* codes.^54^

To estimate the binding free energies of the protein/DNA complex in the different poses and in presence of PTMs we have used the MM/GBSA^55^ approach as available in the AMBER22 package.^56^ The input parameters were kept as such (internal and external dielectric constants of 1 and 78.5, respectively), and the energy decomposition by residue was turned on.

## Author Contributions

The article was planned and realized with contributions by all the authors. All the authors approved the final version of the manuscript.

## Acknowledgements

The authors thank GENCI and Explor computing centers and the Platform P3MB for computational resources. The CNRS and French Ministry of Higher Education and Research (MESR) are also gratefully acknowledged for funding of the GAVO program. is gratefully acknowledged. A.M. acknowledges funding from DISCOVER-UAH-CM project (Ref.: REACT UE-CM2021-01), 282 cofounded by Community of Madrid (CAM) and European Union (EU), through the European 283 Regional Development Fund (ERDF) and supported as part of the EU’s response to COVID-19 pandemic.

