## Supplementary Information for "How SARS-CoV-2 alters the regulation of gene expression in infected cells^†^"

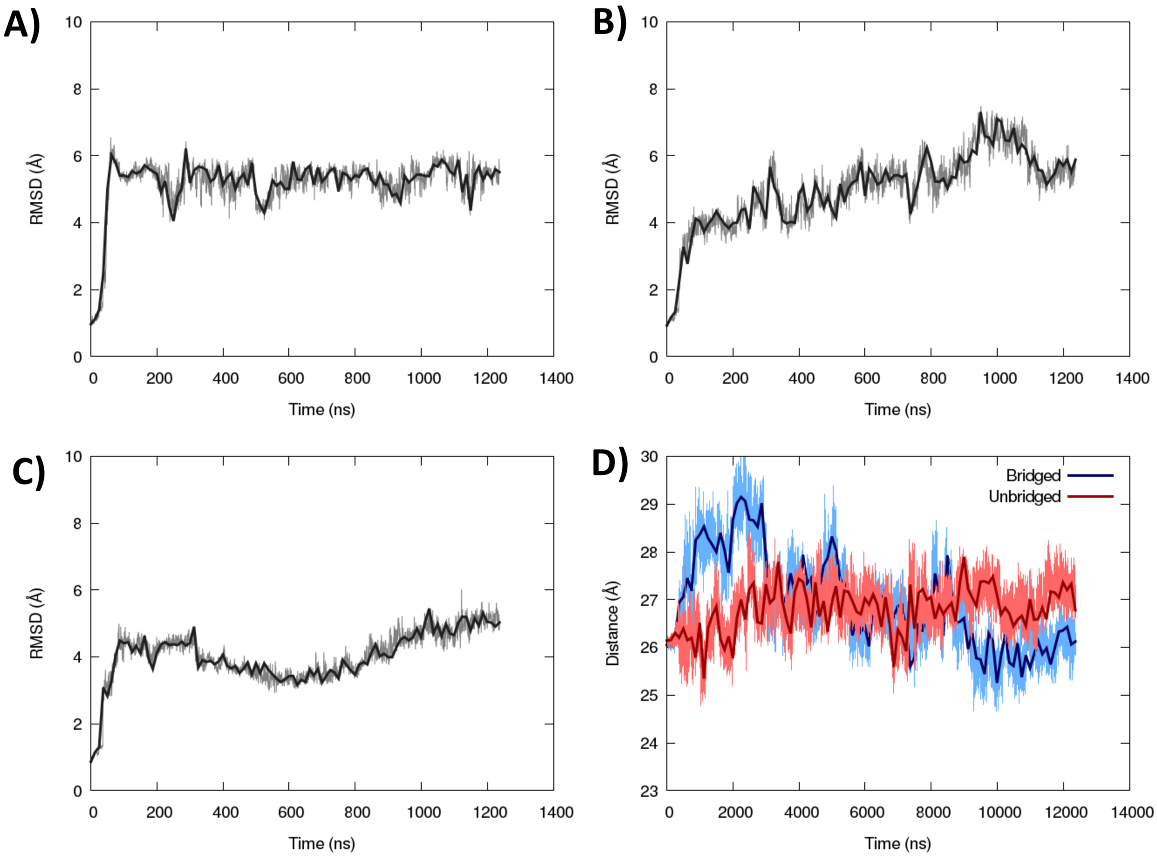


Figure S1. Time series of the RMSD for apo ORF8 in monomeric form (A) in the sulphur-bridged dimer (B) and in the non-sulphur-bridged dimer (C). Time evolution of the distance of the center of mass between the two monomers of ORF8 for the bridged and non-bridged states (D).

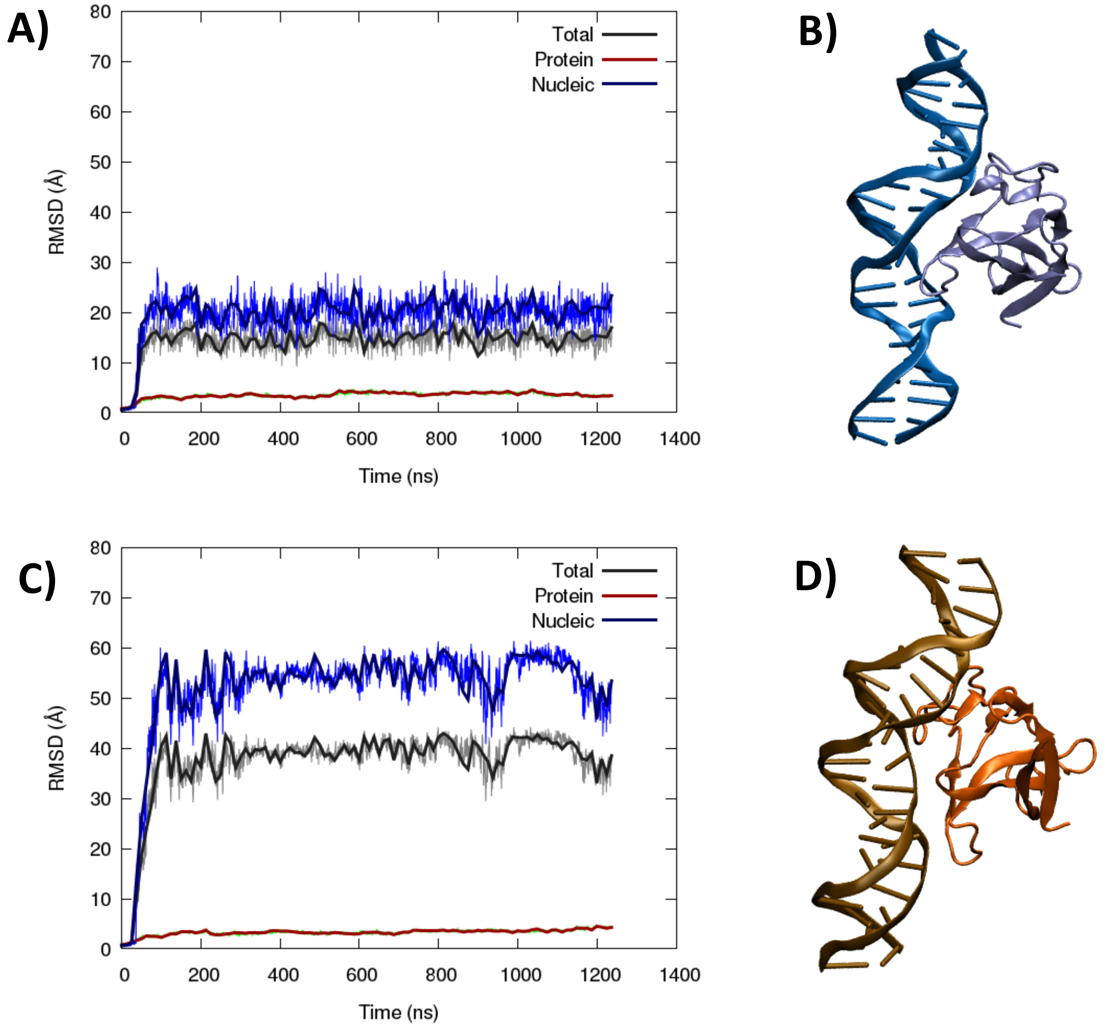


Figure S2. Time evolution of the RMSD for the DNA-bound monomeric ORF8 in the major groove (A), the initial docking pose is given in panel B). Time evolution of the RMSD for the DNA-bound monomeric ORF8 in the minor groove (C), the initial docking pose is given in panel D).

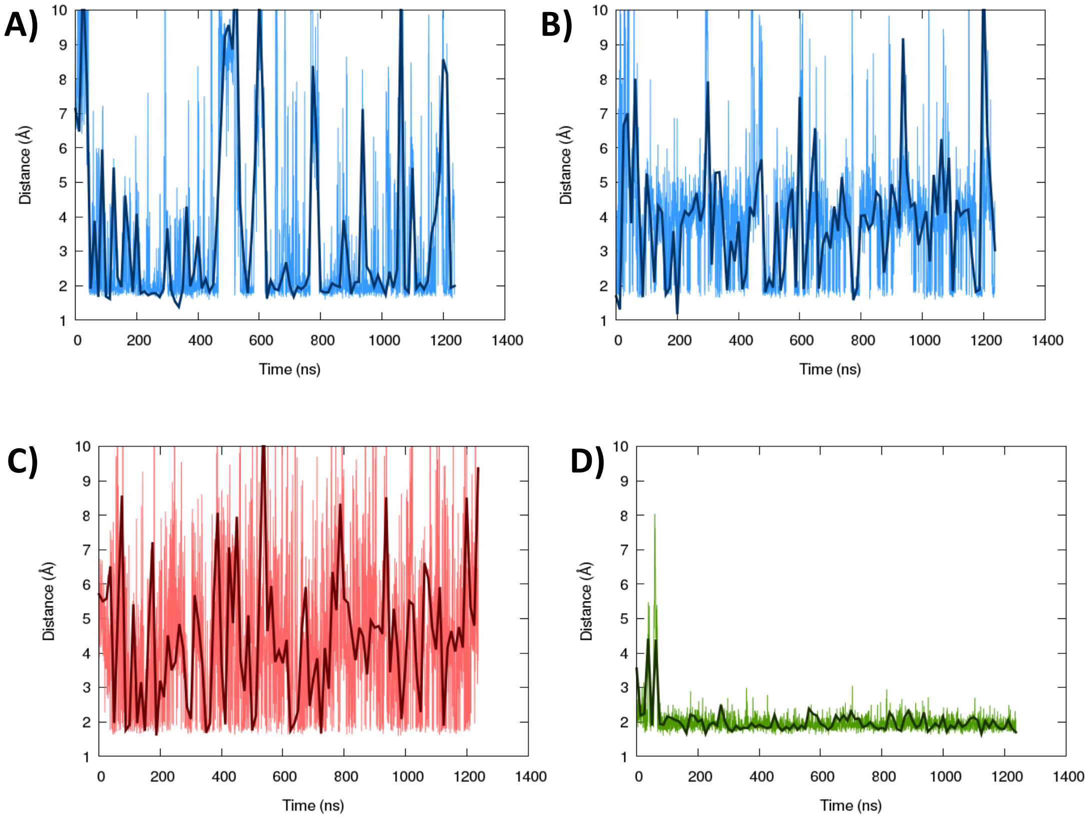


Figure S3. Time evolution of the distances between the R36, K37, and S38 amino acids and the most proximal nucleotides for the major-groove-bound ORF8, i.e. R36/T11 (A), R36/T12 (B), K37/A29 (C), and S38/C31. See Figure 4 in the main text for the corresponding distributions.

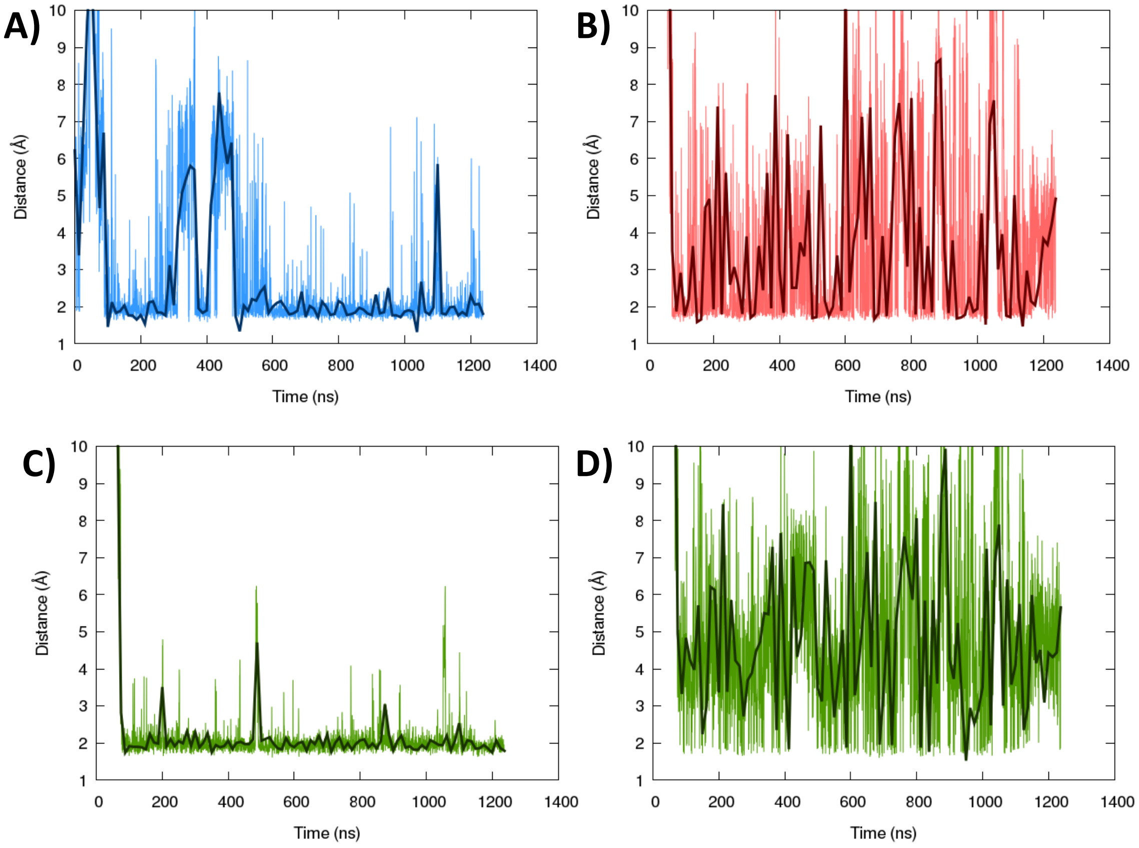


Figure S4. Time evolution of the distances between the R36, K37, and S38 amino acids and the most proximal nucleotides for the minor-groove-bound ORF8, i.e. R36/G40 (A), K37/C42 (B), S38(OH)/T41 (C), and S38(NH)/T41. See Figure 5 in the main text for the corresponding distributions.

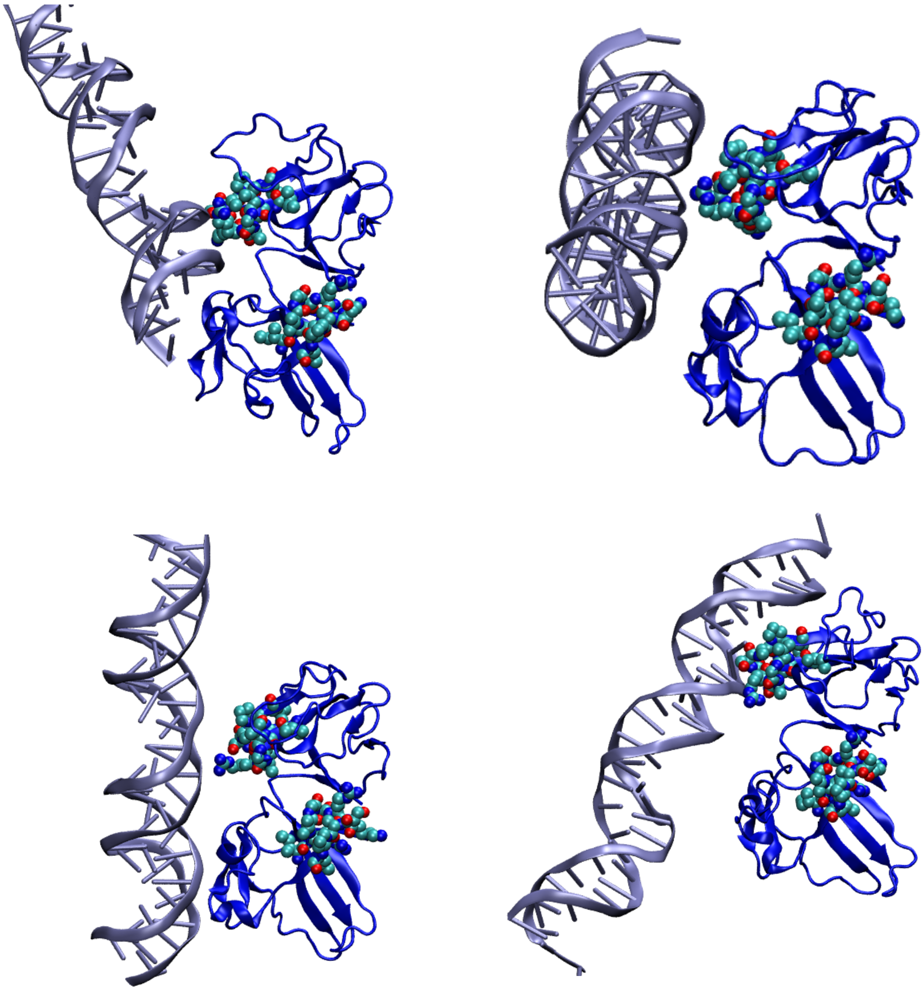

Figure S5. Representative snapshots of the dynamics between the dimeric ORF8 and DNA, showing more labile interactions with the ARKS motif (highlighted in van der Waals representation) and the absence of cooperative binding involving the two monomers.

| **Major Groove** | ΔG (kcal/mol) | **Minor Groove** | ΔG (kcal/mol) |
|---|---|---|---|
| WT | -45.6 ± 8.2 | WT | -28.4 ± 9.5 |
| AC | -39.8 ± 9.8 | AC | -31.4 ± 13.6 |
| Me1 | -47.8 ± 9.1 | Me1 | -55.6 ± 13.9 |
| Me2 | -39.6 ± 6.8 | Me2 | -48.4 ± 15.9 |
| Me3 | -43.7 ± 9.4 | Me3 | -41.4 ± 13.9 |

Table S1. Binding free energy between ORF8 and DNA in the major and minor groove binding modes, estimated with the MM/GBSA approach for the wild type protein and for the systems including PTMs on K37.
